# Extending the Boundaries of Cancer Therapeutic Complexity with Literature Data Mining

**DOI:** 10.1101/2022.05.03.490286

**Authors:** Danna Niezni, Hillel Taub-Tabib, Yuval Harris, Hagit Sason-Bauer, Yakir Amrusi, Dana Azagury, Maytal Avrashami, Shaked Launer-Wachs, Jon Borchardt, M Kusold, Aryeh Tiktinsky, Tom Hope, Yoav Goldberg, Yosi Shamay

**Affiliations:** Faculty of Biomedical Engineering, Technion – Israel Institute of Technology, Haifa, Israel; Allen Institute for AI, Tel Aviv, Israel; Allen Institute for AI, Seattle, USA; Bar-Ilan University, Ramat-Gan, Israel; The Hebrew University, Jerusalem, Israel

## Abstract

Drug combination therapy is a main pillar of cancer therapy but the formation of an effective combinatorial standard of care (SOC) can take many years and its length of development is increasing with complexity of treatment. In this paper, we develop a path to extend the boundaries of complexity in combinatorial cancer treatments using text data mining (TDM). We first use TDM to characterize the current boundaries of cancer treatment complexity and find that the current complexity limit for clinical trials is 6 drugs per plan and for pre-clinical research is 10. We then present a TDM based assistive technology, cancer plan builder (CPB), which we make publicly available and allows experts to create literature-anchored high complexity combination treatment (HCCT) plans of significantly larger size. We develop metrics to evaluate HCCT plans and show that experts using CPB are able to create HCCT plans at much greater speed and quality, compared to experts without CPB. We hope that by releasing CPB we enable more researchers to engage with HCCT planning and demonstrate its clinical efficacy.

## Introduction

### Combination Therapy in Cancer

The number of available cancer treatments is increasing in size and in diversity, involving a wide variety of modalities such as surgery, chemotherapy, targeted therapy, cell therapy, radiation therapy, immunotherapy, etc.^1^ Oncology relies heavily on pharmaceutical intervention and drug combinations are used in most cancers, even though there is a strong incentive to develop monotherapies as a substitute. The most ideal example of such successful transition is the approval of Imatinib monotherapy for chronic myeloid leukemia (CML) and gastrointestinal stromal tumors (GIST) over chemotherapy^2^ On the other hand, for some cancers it seems that the current optimal solution is adding more drugs to the standard of care (SOC) instead of replacing them.

This approach is well demonstrated in the case of Diffuse Large B-cell Lymphoma (DLBCL).^3^ The history of the SOC in DLBCL can teach us a lot on how complex treatments are formed. After testing multiple chemotherapeutic drugs from the 1980s until early 1990s, a four-drug regiment named CHOP (Cyclophosphamide, Hydroxydaunorubicin, vincristine and Prednisone or Prednisolone) was selected as the SOC. In the late 1990s, Rituximab, a CD20 antibody was tested as monotherapy in DLBCL, with the aim to replace CHOP. Even though Rituximab had an impressive activity, it was not superior to CHOP, and eventually was tested in combination.^4^ The 5-drug combo RCHOP was approved in 1999 and in 2015 a kinase inhibitor, Ibrutinib was tested as a monotherapy replacement.^5^ Again, even though it showed activity, it was not superior to the SOC and was tested in combination to form a 6-drug regimen– RCHOP-I. The fact that it took 40 years to optimize this 6-drug regimen as SOC indicates that there is an unmet need to accelerate discoveries of complex drug regiments and treatment plans.

One can view the difference between the treatments of CML and DLBCL as two opposing arms of scientific reasoning: reductionism vs. complexity. Complexity theory describes the behavior of complex systems in multiple domains by studying the relations of multiple elements, opposed to a reduction to the most basic elements. Complex systems consist of multiple parts exhibiting synergistic relations which eventually result in novel emergent properties beyond the properties of individual elements^6^. As mentioned above the increase in treatment complexity can occur via a stepwise addition of drugs, but also through other routes. One such important route is ‘horizontal transfer’ importing a synergistic pair from one cancer to another. For example, the combination of surgery and the adjuvant chemotherapy FOLFOX (oxaliplatin, 5-FU and leucovorin) has been remarkably efficient in colon cancer^7^ and was transferred to many other cancers such as bladder cancer.^8^ In addition, combination of immune checkpoint inhibitors of PDL1 and CTLA4 were shown first synergistic in melanoma and then were transferred to other cancers^9^ such as lung^10^, pancreas^11^, liver^12^ and colon^13^ cancers.

Here we propose to handle the high difficulty of designing useful complex treatment plans by exploiting data-mining over the scientific literature, which was generated by decades of trials and errors of combination therapy, and which is too vast for a clinician to follow. One can imagine the following theoretical thought experiment: if a clinician, for example an oncologist, could read, remember and understand all the papers published in medical history, would they propose a more complex treatment plan? what is the resolution and detail such a plan can contain? and what would be the burden of proof for the new plan? We have developed a protocol and a supporting software application to help the clinician access the relevant literature.

### Text Data Mining (TDM) for High Complexity Combination Therapy (HCCT)

As we’ll show in the following section, researchers are typically very conservative in forming and testing high complexity combination therapies (HCCT), with less than 1% of the drug combinations submitted to clinicaltrials.gov having 5 drugs or more.

Indeed, to motivate the creation of higher complexity treatment plans a researcher needs to consider the interactions between the suggested drugs and the target cancer, as well as the mutual interactions between the drugs themselves. Furthermore, the researcher needs to show that the plan is more promising than the multitude of alternative plans, which is hard for large plans: e.g., by only considering proper subsets of an *n*-drug plan as its alternatives, the researcher is already faced with *2^n-1^* competing plans to evaluate. As we’ll show in Section “Results/Cancer Plan Builder Tool”, as the number of drugs increases, keeping track of all the documented interactions becomes infeasible. We suggest however, that Text Data Mining (TDM) can be used to mitigate these problems by helping the researcher to organize and structure the literature and to surface relevant findings.

TDM has shown consistent improvements in identifying biomedical entities and relations,^14^ and recently it has been applied to the problem of automatically identifying drug combinations.^15^ Despite these advancements, we do not believe that the state of the art in drug combination extraction is at a level which could be used for fully automated construction of treatment plans (see discussion in Section “Methods/Integrating Supervised Machine Learning (ML) Models”). Instead, we present a TDM-assisted planning tool (Cancer Plan Builder, or CPB) that puts the researcher at the center of the planning process: based on initial details of the target cancer, CPB presents the researcher with relevant drugs mined from the literature. The researcher can add selected drugs to a plan, and in turn see suggestions for additional drugs that combine synergistically with the ones already added. At every stage, the researcher can inspect evidence from the literature and use these to assign scores to drug-drug and drug-cancer interactions within the plan. CPB then calculates a global plan score based on these association scores.

CPB allows a researcher to create multi-drug treatment plans based on TDM suggestions and evidence from the literature. The use of TDM reduces planning time and guarantees that the researcher does not overlook important published information. The scoring system allows researchers to compare different plans for the same condition and provides means to prioritize certain plans over others. By releasing CPB we hope to reduce the entry barriers for HCCT, enable more researchers to engage with it and lead to gradual unraveling of its best practices and clinical efficacy.

In the following sections, we: (i) characterize the current and possible landscape of combination therapy in cancer and unravel the boundaries of treatment complexity; (ii) present CPB, a graphical tool based on text mining which assists researchers in creating, comparing and motivating literature backed complex treatment plans; (iii) develop scores and metrics to compare HCCT plans; and (iv) use these metrics to evaluate CPB, showing that experts assisted by CPB can accurately reproduce existing treatment recommendations, as well as create higher complexity plans which are more cohesive and better grounded in the literature, compared to plans generated by experts without CPB. Plan generation time is also considered, showing remarkable time savings: 30-60min for plan generation with the CPB compared to multiple work days for plans generated without it. The tool is publicly accessible here: https://planbuilder.apps.allenai.org/

## Results

### Characterizing the current complexity of cancer treatments

Before exploring the generation of complex treatment plans, we sought to characterize and analyze the status of drug combination therapy in research and clinical trials. We used two different methods to capture the number of pharmaceutical interventions applied in different cancers to quantitively characterize the level of complexity in combination therapy (**Figure 1a**). We mined drug combinations in research from PubMed abstracts and clinicaltrials.gov for clinical stage combinations. Our text mining approach used both explicit and implicit drug combinations. In the explicit search, we search for mentions of specific drug names in the described treatment (e.g., for two drugs, “a combination of dolutegravir and lamivudine was administered …”). In the implicit searches, we search for mentions of the number of drugs (e.g., “the 2-drug regimen proved effective …”). To accurately capture the syntax of implicit combinations, we used the SPIKE engine for syntactic structure search^16^ and ranked the frequency of implicit drug combination phrases (**Supplementary Fig S1**). The most frequent phrasings were ‘X-drug combination’ and ‘X-drug regimen’. When the two approaches were plotted, it was clearly evident that there is a logarithmic decline in frequency (string negative correlation) of each additional drug to the combination, ending at 6 drugs when the drugs are mentioned explicitly by name (**Fig 1b**). Additionally, we found that when drugs are explicit, the number of drugs in the combination drops down in the exact same manner in both research and clinical trials **(Figure 1b**). But when the drugs are implicit, the decline from two to four drugs is moderate. This can be explained by PubMed abstract’s word limit, where describing a 4-6 drug combination by the drug names will use valuable word space. Furthermore, there are no more than six drugs mentioned explicitly in any combination, while when the drugs are implicit, combinations of up to 10 drugs arise from the search. Additionally, we can learn that combinations in clinical trials are mostly studied with 2-3 drugs while combinations of 4 drugs represents less than 10% and 5 drugs less than 1% of clinical trials. The feasible limit of drug cocktails has only a handful of clinical trials with 6 drugs per treatment. Though it is possible that there are clinical trials with more than 6 drugs that were missed due to the explicit search, it is not likely to post a clinical trial without specifying the drugs especially in clinical trials. An additional search for sizes of implicit drug combinations not constrained to cancers type (“Pubmed ES”) revealed that there are >50 fold more publications in the >6-drug combination category in other diseases, mainly to anti-HIV cocktails related research as well as antibiotic cocktails (**Fig 1b**). Next, we wanted to quantitively characterize the drug combination space and its relation to the size of the treatment plan and the number of possible drugs to choose from for each cancer type. Our main motivation was to understand how much of the possible combination space is covered in the scientific literature and how much of it can be tested and validated experimentally. To calculate the number of possible drug combinations we used **eq. 1**, an equation for calculating the number of possible ways to select groups of ‘r’ drugs out of ‘n’ possible drugs, with no repetitions in the same combination. For example, there are 190 possible 2-drug combinations for a cancer type with 20 optional drugs to choose from, but for a cancer with 40 available drugs the number of possible 2-drug plans is 780. As expected, there is a rapid growth of the complexity, a combinatorial explosion with increasing dependency on the numbers of drugs per plan **(Fig 1c).** We show that even with automated in vitro experiments which can handle 10^5^ combinations it is impossible to cover more than 3-drugs per plan. To characterize the combination possibility space for each cancer type, it is required to know the number of possible drugs that are relevant for that particular cancer. To this end, we used the SPIKE search engine to quantify the number of drugs studied in each cancer by capturing sentence-level co-occurrence of drug name and cancer name. We found marked differences between the cancer types (**Figure 1d**). Breast cancer for example is the most studied cancer, with more than 400 possible drugs, as opposed to rhabdomyosarcoma for example, which has less than 20 possible drugs to choose from. Thus, with this approach the number of possible plans for each cancer type is directly correlated to the volume of research performed on that cancer.

**Figure 1.**
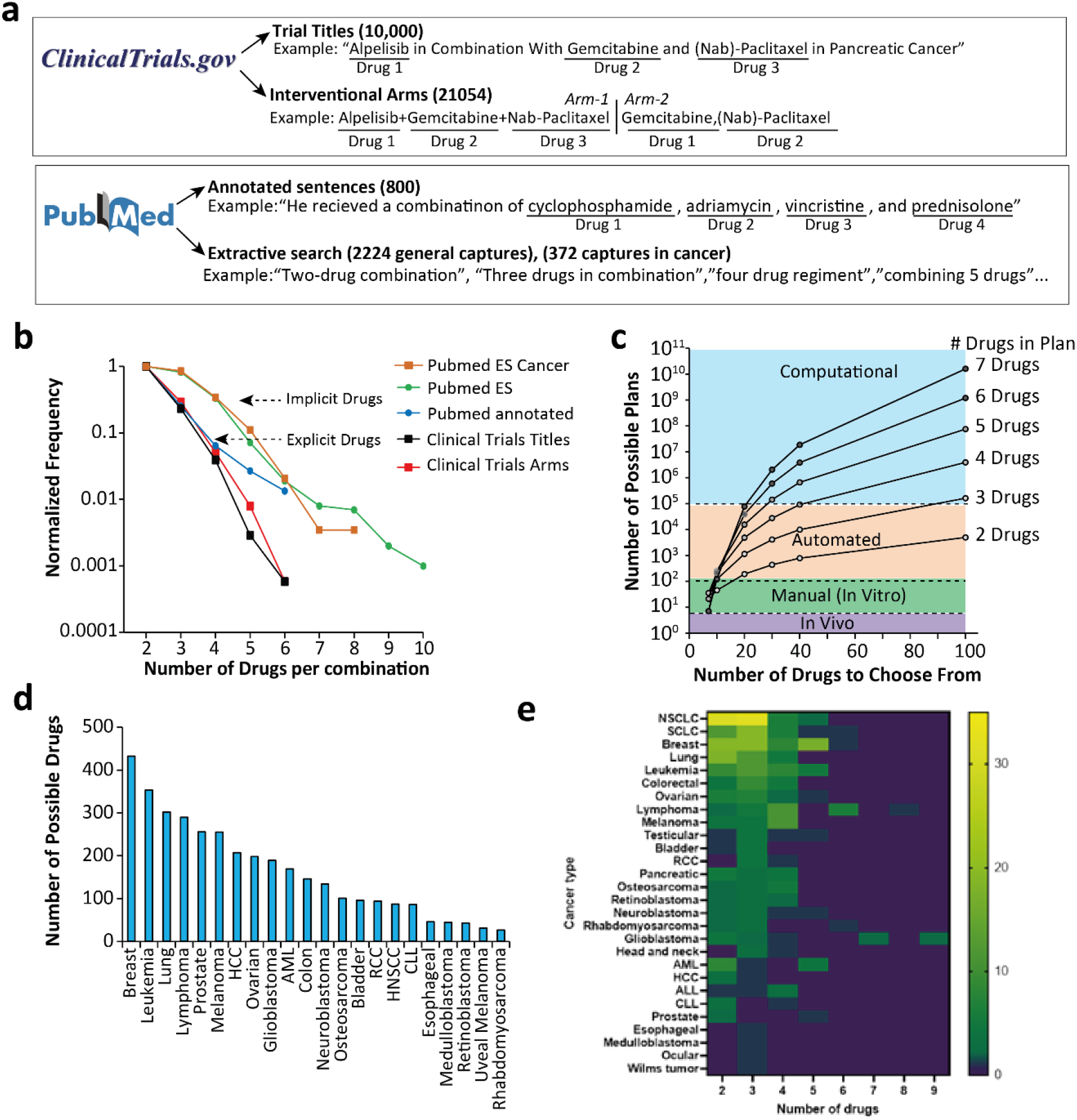
The current and possible landscape of combination therapy in cancer. **a)** approaches to characterize combination therapy from clinicaltrials.gov (top) or PubMed (bottom). **b)** Normalized frequency of drug combinations in treatments for cancers derived from clinialtrial.gov, PubMed and manual annotations. **c)** Number of possible plans according to the number of possible drugs in plan and number of drugs to choose from. **d)** Number of possible drugs per cancer type, extracted from PubMed abstracts via SPIKE. **e)** Frequency of implicit drug combinations sorted by cancer type, extracted from PubMed via SPIKE.

**Eq. 1**-the number of possible ‘r-drug’ combinations for a cancer type with ‘n’ studied drugs.

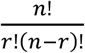

To further evaluate the distribution of treatment complexity in cancer research, we sought to interrogate drug combination distributions within specific cancer types in the scientific literature. We captured the number of implicit combinations (such as ‘five drug regimens’) in PubMed and sorted them by cancer type (**Fig 1e**). Most cancer types had combination of up to four drugs per treatment, and it was quite uncommon to see six or more drugs combined. Lung cancer, breast cancer, leukemia and colorectal cancer don’t have treatments that combine more than five drugs. There might be some bias against combinations of two drugs as many studies use the explicit drug names. This can be seen in some cancers like lymphoma and melanoma where the four-drug combination is more common than combinations of two or three drugs. The upper limit for drug combination, was identified in glioblastoma (GBM) with a combination of nine drugs. This shows that the five- or six-drug limit in clinical trials is being pushed in pre-clinical research. We can speculate that as GBM has poor prognosis, multiple drug combinations are more likely to be tested than cancers that reached a good prognosis with the current SOC. As was mentioned in the introduction, we can see 6-drug combos in clinical trials for DLBCL which took over 3 decades to develop.

## Cancer Plan Builder Tool (CPB)

Based on the current experience with HCCT as described in the Introduction section, we argue that a workflow for generating plans can be derived from the following assumptions:

1. Drugs can be ranked and suggested to the user from the scientific literature based on their connection to an input disease name, associated genes/proteins and additional keywords. This is inspired by the development of the RCHOP-I combination where Rituximab and Ibrutinib were added to the SOC after being first suggested as monotherapies for DLBCL^17^.
2. The user can choose the most relevant drugs based on both the provided score and the highlighted evidence from the literature.
3. Addition of drugs to the plan can be made in a stepwise manner based on drug synergy evidence between all the drugs in the plan and not necessarily based on disease relevance. This is inspired by cases like that of PDL1 blocker which was shown to be synergistic with CTLA4 blocker in melanoma^13^ and then added to other regimens involving CTLA4 for other cancer types^9^.

With these assumptions in mind, planners could in principle employ a search tool like PubMed to search for papers with mesh headings relating to the disease conditions and to therapeutic agents. Planners could then screen the results to create a short list of first line treatments for the cancer, followed by additional PubMed queries to obtain papers which reference these agents, and screening them for additional agents which synergistically combine with the existing ones. This process can then proceed in an iterative fashion, continuously adding treatments which are synergistic to existing ones.

While this workflow is feasible, the need to screen the returned abstracts at each stage of the process does not allow treatment plans to scale beyond a handful of treatments. For example, an initial PubMed search for “(Melanoma[MeSH Terms]) AND (therapeutic uses[MeSH Terms])” returns 14459 results. Thus, the screening required to establish a comprehensive list of promising first line treatments may already be prohibitively time consuming, even without considering additional searches which could identify synergistic treatments.

To address these limitations, we designed and implemented a dedicated cancer plan builder application (CPB), which streamlines the above workflow and dramatically reduces the screening time. The time saving is achieved by allowing to search for co-occurrences of treatments with cancer types, genes, and other treatments, displaying this information in an organized manner, allowing review and acceptance of suggested treatments, and automatically refining the queries as new treatments are added to the combination. Treatment planning with the app proceeds as follows: (**Fig 2a, 2b,** see also “**Methods/CPB tool development**” for details of the extraction methods used)- (i) the planner inputs the cancer type, the patient genetic history and biopsy findings, and optionally other keywords of value (e.g., metastasis if the cancer is at that advanced stage, etc.) (ii) the app performs an initial search and extract and displays a list of treatments ranked by their relevance to the input conditions (up to 100 treatments). Each treatment is associated with evidence from the literature showing its relation to the input conditions. (iii) the planner reviews the treatment and evidence and moves one or more treatments into a planning canvas showing the current plan (iv) the planner can click any treatment on the canvas to trigger a search for treatments which combine with it synergistically (v) as before, the planner sees the evidence for the pairwise synergy before deciding whether to add a synergistic treatment to the plan (vi) the planner can connect an added treatment to other treatments in the plan and inspect evidence for potential synergies (or adverse reactions) further increasing the cohesiveness of the plan. (vii) CPB assigns a default association score for each connection and this score can be modified by the planner based on inspected evidence (default association scores are based on treatment-condition and treatment-treatment co-occurrence metrics, see details in the section **Methods/CPB Tool Development**). These pairwise scores are then used by CPB to assign global scores to plans, and these can in turn be used to prioritize and rank plans (see section “**Evaluating Treatment Planning with CPB**”). The process continues in an iterative fashion where the planner can experiment with adding and removing treatments based on the evidence.

**Figure 2.**
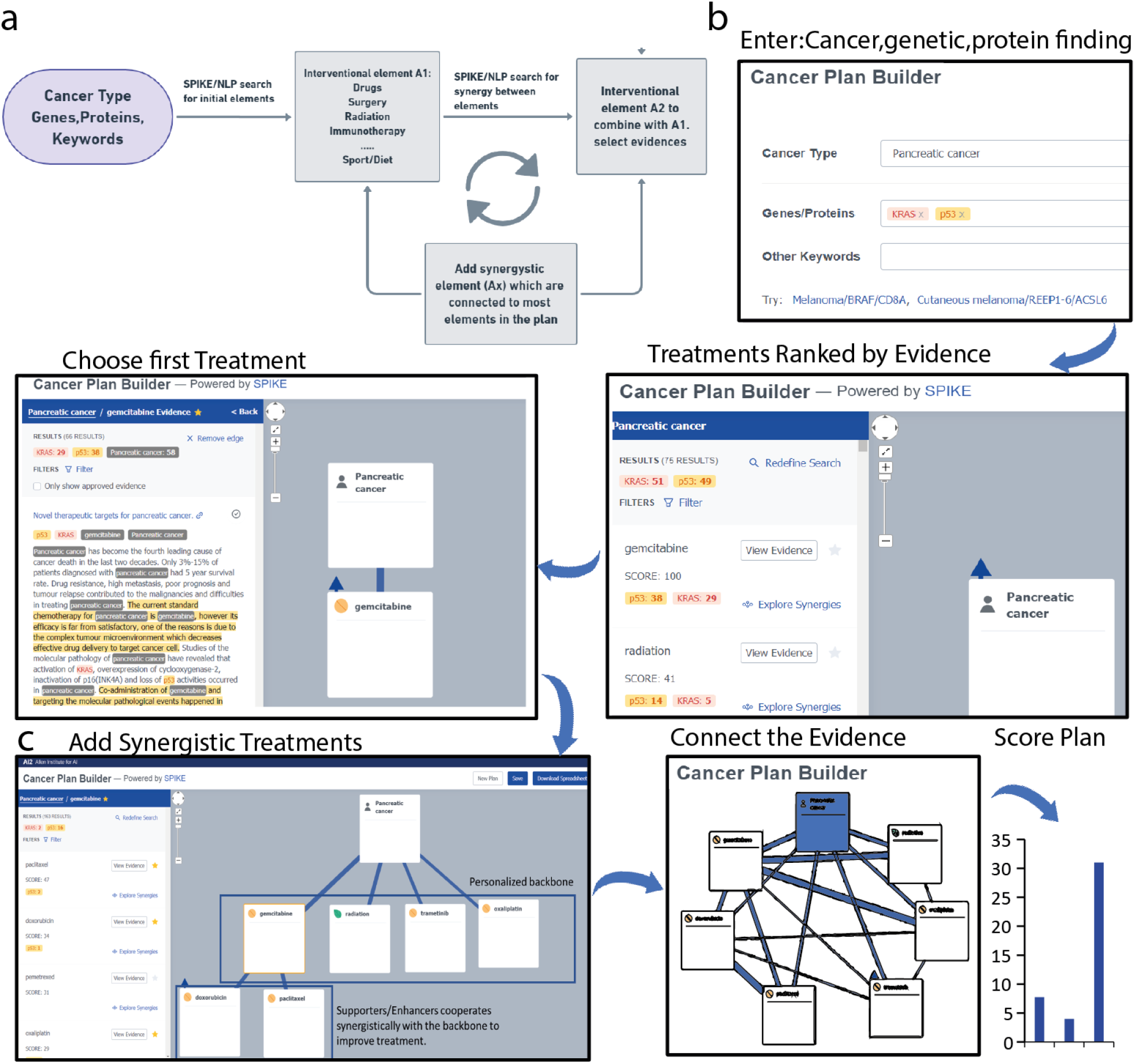
Workflow of building a plan with CPB. **a)** thought process and fallback strategy. **b)** an example of the process and its result, the user built the personalized backbone treatment with drugs that are directly associated with pancreatic cancer, added the supporting drugs and finally, found the interconnectivity and exported the score of the plan.

Note that in contrast to using a standard search engine like PubMed, planners don’t need to screen large lists of papers in order to identify candidate treatments. The app automatically extracts treatments and ranks them by the strength of their association to the input conditions (for first line treatments) or to other treatments which are already in the plan (for additional treatments. See the Methods section for the details of this process). Planners still need to validate short evidence snippets associated with the treatments, but as we’ll demonstrate, this process is an order of magnitude faster than traditional screening and allows plans to scale to a significantly larger number of components while staying reasonable and consistent.

Finally, CPB offers a number of additional utility functions, not involving text mining. Users can add comments to treatments or connections in the plan (e.g., to document timing and dosage information), add treatments not suggested by the CPB, save and share plans with other users, etc. The CPB tool can be found here: https://planbuilder.apps.allenai.org/

### Evaluation

To evaluate our method, we start by comparing the individual drugs that were highly ranked by CPB to the standard of care (SOC) in 12 different cancers, showing that the vast majority of SOC treatments are correctly identified and suggested by CPB, and that CPB suggests many additional relevant treatments. We then turn to evaluate end-to-end treatment planning with CPB. We start by defining evaluation metrics, establishing baselines and continue to analyze and compare treatment planning conducted by experts using CPB to treatment planning done with traditional literature search, without CPB.

### Evaluating the Ranking of CPB Drug Suggestions

To evaluate the relevance of the app drug suggestions, we measured their overlap with the standard of care (first line therapy) as reported by cancer.org and UpToDate. We considered any interventional treatment in the SOC including non-pharmacological interventions such as radiation, surgery etc. In addition, we looked at the number of FDA approved treatments for a certain indication, beyond the standard of care such as second/third-line therapy and advanced investigational treatments. The app ranks up to 100 drugs per case and thus we looked at the top 10 drugs as well as total 100. We conducted two experiments, each involving 12 cancer cases: 6 of which were general cancer types and 6 were cases of personalized cancer treatment, where the cancer is associated with a known targetable mutation present in a patient. In the first experiment we found that in all 12 cases the vast majority of SOC drugs were also suggested by CPB according to both UpToDate and cancer.org (**Fig 3a**). We found that though there are differences in the SOC reported in the two resources, there is above 85% coverage of the app between them for all cancers. There was 100% coverage for 9 out of 12 (75%) cases according to cancer.org and 8 of 12 (67%) for UpToDate. The non-100% coverage cases where all above 80% except for AML and liver cancer. Examining treatments which were not suggested by the app like ‘bone marrow transplant’ and ‘CAR T cells’ for FLT3 mutated AML and Transarterial chemoembolization (TACE) for liver cancer, we learned that the app is highly compatible with simple pharmacological treatments and standard drug entities but not with sophisticated procedures, or novel treatment classes such as biological cell therapy (**Fig 3a**).

**Figure 3.**
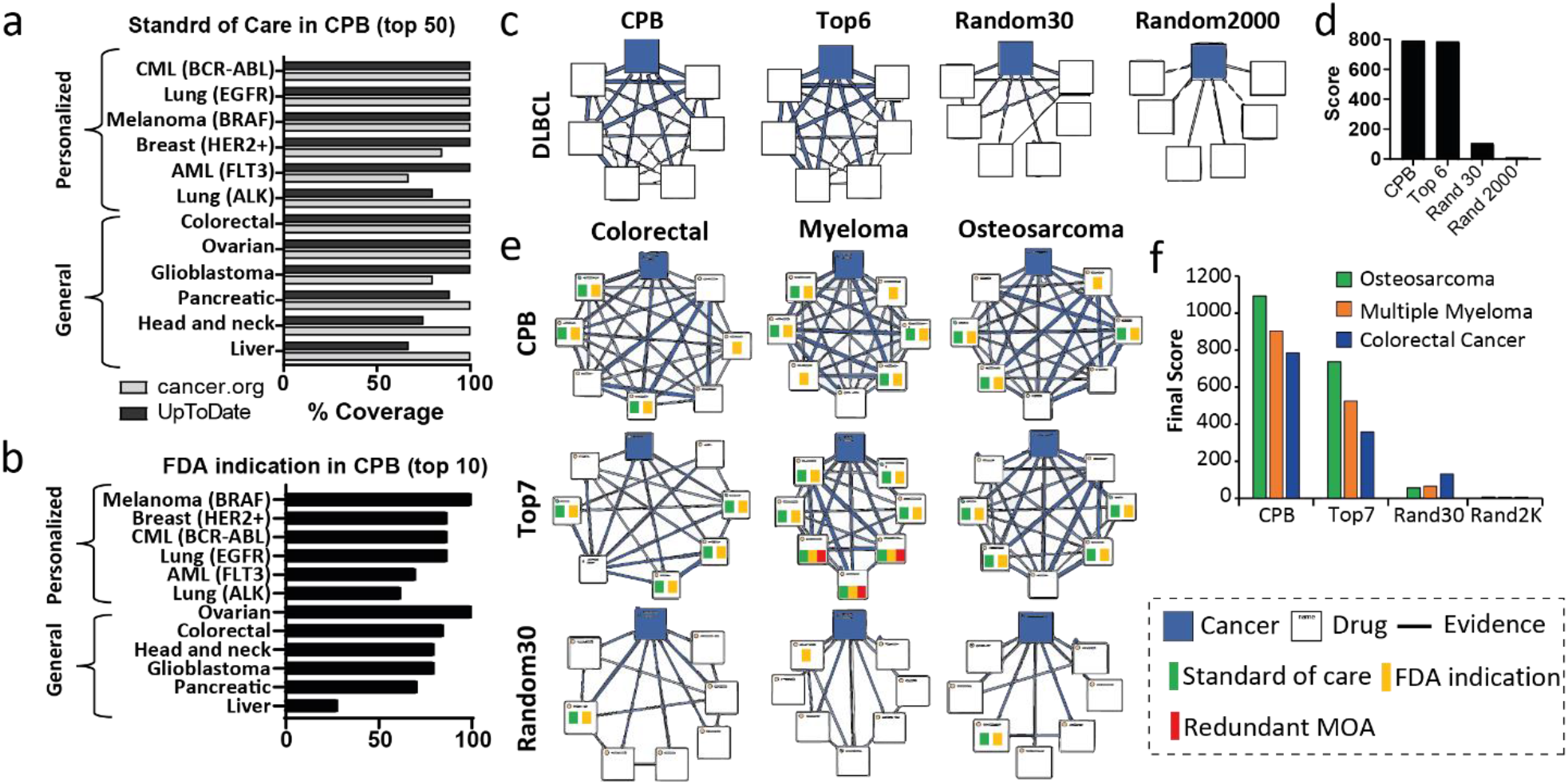
Evaluation of CPB app. **a)** % of Standard of care treatments in the top 100 ranked by the app as covered in uptodate.com or cancer.org. **b)** % of FDA approved drug for the specified indication in the top 10 ranked drugs by the app. **c**) 6 element plans for DLBCL. All plans were generated with the app either with the SOC, drugs chosen from the top 7 ranked drugs, or random pick from top 30 or 2000 FDA approved drugs. **d)** plan evaluation score using with (weights/nodes)×edges. **e)** 7 element treatment plans for 3 cancer types (Colorectal, Multiple Myeloma and Osteosarcoma). All plans were generated with the app either with a human choosing drugs based on evidence, drugs chosen from the top 7 ranked drugs, or random pick from top 30. The drugs were verified for correct FDA indication approval, standard of care and mechanism of action (MOA). **f)** plan evaluation score with (weights/nodes)×edges

In the second experiment, we wanted to measure how many of the drugs suggested by CPB are generally relevant for the cases in question, irrespective of whether they are a part of the first line treatment. To approximate this, we measured how many drugs in CPB’s top 10 suggestions are FDA approved for the relevant indication and found that indeed, more than 75% of top 10 suggested drugs approved for the relevant cancer type (**Fig 3b**). Interestingly, for liver cancer, only 25% of drugs were FDA approved, which can be explained by the lack of approved options for hepatocellular carcinoma, the most common cancer of the liver^18^. When comparing personalized and general cancer types, there are more FDA approved drugs in the personalized cases (82.5% vs. 74.2%, respectively), while there is no significant difference between the two in the standard of care (80% for both) This data supports our assumption that including case related keywords, can increase the quality of drug suggestions which facilitates generation of reasonable plans.

### Evaluating Treatment Planning with CPB, Method and Baselines

#### Evaluation Metric

To evaluate treatment planning with CPB we start by defining evaluation metrics and establishing a baseline. Our general premise is that other things being equal a better HCCT plan is a more cohesive one: that is, it should exhibit high synergy between the treatments involved. To formalize this we represent a plan as a weighted undirected graph *G* = (*V, E, w*) with a set of nodes *V* (the treatments, plus an additional node representing the input condition), a set of edges 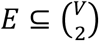 (edges between pairs of elements from *V*) and a weight function 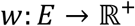 assigning an association score to connected treatments, or to a treatment connected to the input condition. For *e* = (*v*1, *v*2) ∈ *E, w*(*e*) is set by CPB to an initial value based on the number of times *v1* and *v2* appeared together in PubMed papers, their proximity in these papers as well as other factors (see details in the Methods section). This initial score is designed to approximate the quality of the evidence indicating synergy between *v1* and *v2*, but it can be overridden by the expert using CPB. An expert could down-weight cases where the drugs are mentioned frequently but in a negative or otherwise controversial circumstances (e.g., the most frequent drug appearing in papers together with COVID-19 is hydroxochlorquine, but the efficacy of the drug is not generally accepted). An expert could up-weight an edge for edges involving treatments which are promising but very new, as these may be underrepresented in the current literature.

Based on this formulation, we defined several relevant parameters that could be comparable between different plans. We consider the level of confidence *P* = ∑_*e*∈*E*_ *w*(*e*) (the sum of weights in the graph) as the cumulative quality of evidence in the plan. The number of elements in the plan, *N* = |*V*|, can be regarded as the plan complexity index. The number of edges, *R* = |*E*|, can be regarded as the level of inner-synergism within the treatments. A strong plan is likely to have many interconnections between the different nodes (interventions in the plan). This will probably result in high R/N which, when multiplied by the amount of information (P) would lead to the total score reflecting the importance of these variables. Thus, our score is defined as (R/N)×P or (edges/nodes)×weights (**Table 1)**. We note that selecting this score as our evaluation metric is somewhat arbitrary, and that viable alternatives do exist. The score also relies heavily on edge weights, which as discussed above, can be sometimes misleading (or if adjusted manually, not fully objective). Nonetheless, we believe that this simple score reflects the intuition of cohesion through high inter-connectedness and that in practice, as long as users don’t try to explicitly optimize for it, it can be used to prioritize cohesive plans over less cohesive ones. Indeed, while planners can see and tweak pairwise association scores while planning, global scores are not visible while planning and we instruct planners to only trigger their calculation once plans are completed for comparison purposes.

**Table 1.**

| symbol | represents | function |
| --- | --- | --- |
| P | sum evidence | amount of information |
| R | sum relations | Connectivity variable |
| N | sum nodes | Connectivity variable |
| $P * (R/N)$ | multiplication | Final Score |

#### Evaluation Baselines

Based on the proposed scoring system, we tested the generation of four 6-element plans for DLBCL, our inspiration for the project (**Fig 3c**). We compared a plan with the SOC manually inserted in the app, to an automated plan where the first top 6 drugs suggested by CPB are taken together as a single plan. For further comparison, we generated one plan which included 6 random drugs from top 30 suggestions (Rand30) and one with 6 random drugs from a list of 2000 FDA approved drugs (Rand2000). The results clearly show that the plan with the SOC and the top-6 plan were identical and received identical score that was more than 8-fold higher than Rand30 control and orders of magnitudes higher than Rand2000 (**Fig 3d**). As expected, the 6-drug plans consisted of RCHOPI, which showed the best survival benefit in clinical trials^19^, received the highest evaluation score in the app.

We then sought to extend the boundaries of 6-drug combination and generated plans with 7 elements for three case studies and evaluated their score accordingly (**Fig 3e-f**). We chose one adenocarcinoma (colorectal cancer), one sarcoma (osteosarcoma) and one hematological cancer (multiple myeloma). As expected, plans designed by choosing seven random drugs (Rand2000) resulted in scores close to 0, even plans with seven random drugs chosen from top 30 treatments (Rand30) received relatively low scores (<100). Differently from the DLBCL case, the highest score in this evaluation was for the plans generated by the graduate student using CPB, followed by the top7 suggestion (**Fig 3f**). We noticed that both human and top7 scores for the plans for osteosarcoma were higher than multiple myeloma which were higher than colorectal cancer. This observation might reflect the nature of the disease and treatment.

### Evaluating CPB Assisted Treatment Planning

To further evaluate the effectiveness of the app, we compared plans constructed with the help of the app to plans constructed without it. Literature backed treatment planning without the app is a lengthy process so to obtain a collection of treatment plans we devised a planning task and assigned it to biomedical engineering (BME) students as part of a final project in their graduate course. We will first describe this task and its outputs and then report on the comparison results.

### biomedical engineering Course #336041: Advanced Methods in Treating Cancer

We generated fictional cancer patient case studies that were given to 40 graduate students as part of a final assignment in a graduate course titled ‘advanced methods in treating cancer’. The students graduated from biomedical engineering and were either in PhD or MSc tracks in the faculty of biomedical engineering in the Technion, Israel and have background in physiology, anatomy, biology, pharmacology and more. During the semester the students learnt about contemporary methods to treat cancer including: radiotherapy, chemotherapy, immunotherapy (CAR T cells and checkpoint inhibitors) targeted therapy and precision medicine, nanomedicine, oncolytic viruses and anti-angiogenic therapy. In addition, background on hallmarks of cancer and tumor biology were taught as an introduction.

The cases included the patient’s cancer type, genetic finding, protein findings and more personalized characteristics. The task was to generate a complex treatment plan comprising of a minimum of 6 drugs or additional support from any type of non-pharmaceutical interventions such as radiotherapy. As part of the task students were instructed to use standard literature search methods to the best of their ability, but **they did not have access to CPB**. The timeframe for task completion was one month and at the end, we received 23 different treatment plans, each plan is a personalized and complex treatment (**Fig 4a**). We show an example summary of a student project outcome in Ewing’s Sarcoma in **Supplementary Fig S2**. We analyzed the projects and compared the students’ plans with known possible treatments for the same disease from two other resources, Wikipedia.com and the cancer.org. These resources contain summarized common treatment options in a clear and accessible way. As seen in **Fig 4b and supplementary Fig S3**, in all cases, the students offered more complex plans, with more diverse types of elements. The complexity is expressed mostly by incorporating more drugs and wider use of immunotherapy in the treatments. Also, the students included integrative medicine and chronotherapy that are usually not included in standard treatments.

**Figure 4.**
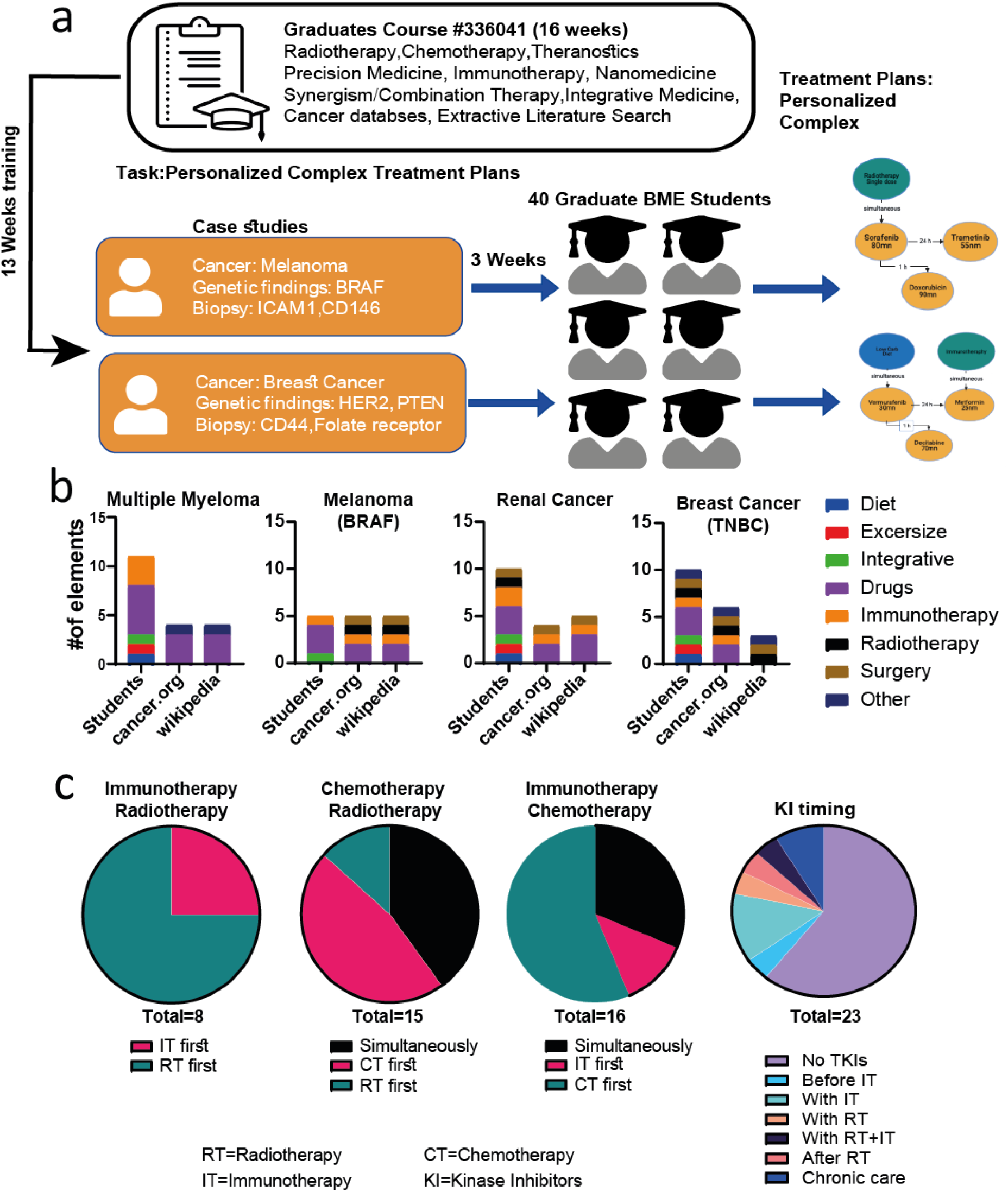
BME Course #336041 Advanced Methods in Treating Cancer. **a)** course workflow. **b)** number of treating elements in constructed plans and the standards of care according to cancer.org and Wikipedia.com, divided by categories. **c)** order of treatments and timing of kinase inhibitors according to the constructed plans.

When analyzing the students’ plans, we searched for an agreement between the students regarding the timing element in their plans (**Fig 4c**). For example, we saw that in the majority of the plans (56.25%) the students suggested to first use chemotherapy and then to apply immunotherapy. This is reasonable, since some chemotherapies cause immunogenic cell death, which can educate the immune system to attack the dead cancer cells, resulting in a more efficient attack on living cancer cells following the immunotherapy^20^. The chronology of radiotherapy and immunotherapy was also widely agreed upon, the majority (75%) suggested the use of radiotherapy first, this is very reasonable biologically since radiotherapy induces an inflammation reaction that will be enhanced later by the immunotherapy and is more likely to reach improved outcomes^21^. Most of the plans concluded that chemotherapy should be used before (46.67%) or at the same time (40%) as radiotherapy. Finally, we noticed most of the plans (60.87%) did not include specific tyrosine kinase inhibitors (TKIs), from those who did include TKIs in their treatment plan the main conclusion was to prescribe them with either radiotherapy or immunotherapy (21.72% from all plans).

### Comparing CPB assisted Plans with Plans Generated by the Students

Next, we selected eight plans from the ones generated by the students and had another student undergo CPB training and use it to generate equivalent plans for the same conditions. Generating plans with the help of CPB was remarkably faster than without, with all CPB plans being generated in under 60 minutes per plan as compared to multiple workdays without it. Then, we compared plans generated using CPB to plans generated without and to the approved treatment plans from Wikipedia and cancer.org (**Fig 5a**). Compared to plans generated without CPB, plans where CPB was used had a significant increase in the number of elements by a factor of 2.5. The CPB assisted plans were versatile and most (62.5%) consist of three types of drugs-tyrosine kinase inhibitors, chemotherapies, and monoclonal antibodies-while the plans generated without CPB consisted mostly of two out of the three elements (**Fig 5b**).

**Figure 5.**
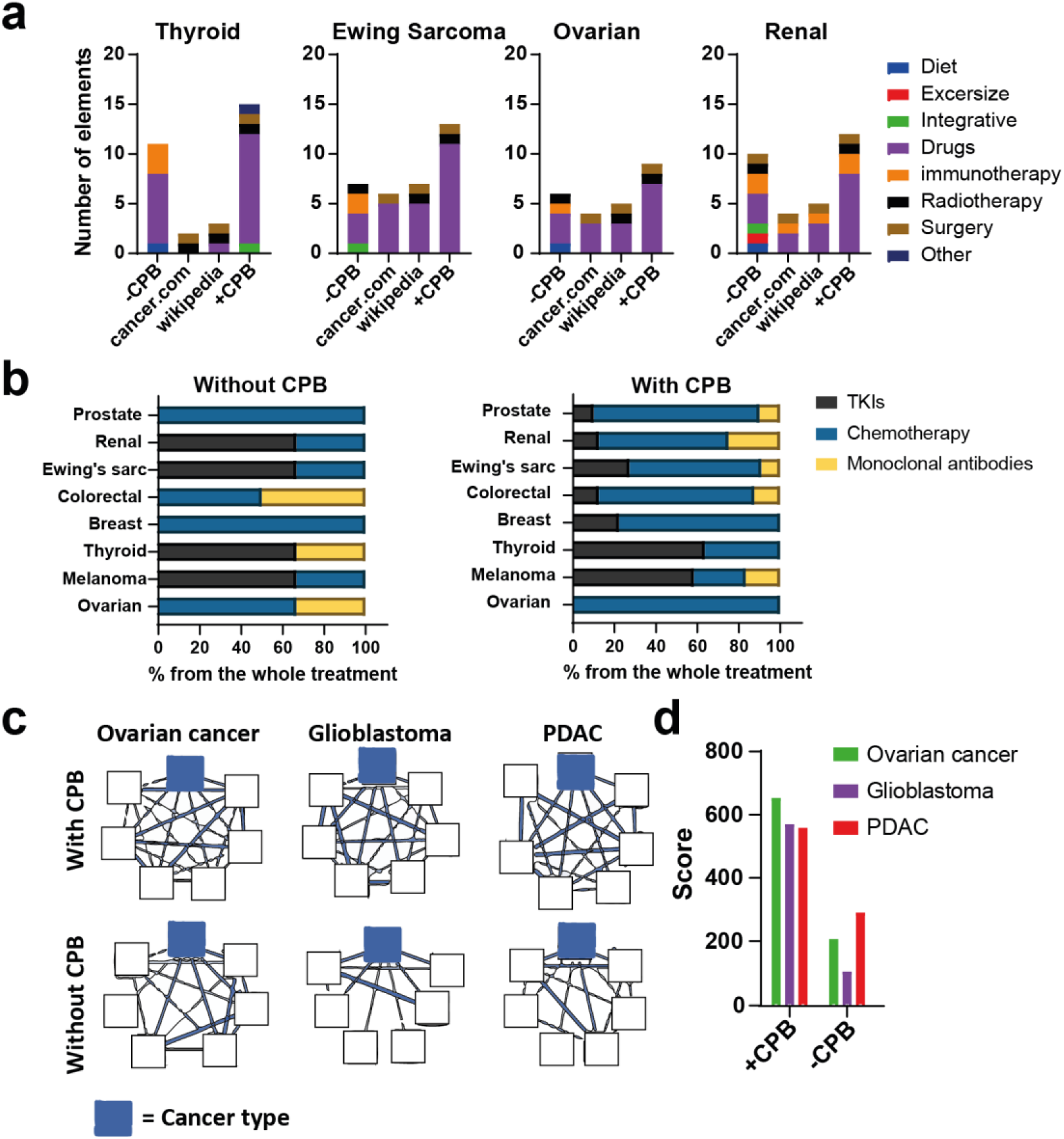
comparison between plans with and without the CPB app. a) number of treatment elements in constructed plans, divided by categories. b) distribution of drug types in plans constructed with and without CPB. c) 6-element treatment plans for four cancer types (DLBCL, ovarian cancer, glioblastoma and PDAC),. All plans were generated with the app either by choosing from the top 6 ranked drugs (CPB) or from the students’ projects. d) plan evaluation score.

In order to further compare the CPB assisted plans with the plans generated without CPB, we manually imported 3 plans generated without CPB to the app for the purpose of scoring them with CPB’s scoring system. In all 3 cases the plans generated with the help of CPB achieved a significantly higher score, indicating higher inner-synergism and greater plan cohesion (**Fig 5c, 5d**).

## Discussion

### Burden of proof in clinical validation of HCCT

In developing CPB, we were inspired by the historic development of multi component treatment regiments with high complexity, such as R-CHOP-I treatment for DLBCL^22^. Our interpretation of the process suggested that monotherapies which fail clinical trials but have significant efficacy can be added in a stepwise manner to the standard of care as a new combination therapy. One of the reasons of the slow development of HCCT is the high burden of proof. The transition from an existing combination to an even slightly more complex one is not small nor simple. By current practice, the burden of proof increases dramatically with increasing complexity, which can be intimidating and highly limiting. For example, if one suggests a novel four-drug combination, it would be required to show it is superior to any drug alone and pair and trio combo which yield 15 different control arms to test one combination hypothesis. It is therefore highly unfeasible to test all possible controls to prove that a complex combination is the optimal therapy.

Instead, we propose a shift in perspective, encouraging the development of higher complexity treatment plans motivated by principles of synergy and cohesion. The use of TDM allows to surface relevant treatment interactions and to differentiate between more promising and less promising plans. When validating the pre-clinical efficacy of HCCT plans, we advocate for a more lenient approach, where a combination is compared to a selected subset of promising alternatives and naturally, to the current SOC. Conducting pre-clinical validation along these lines is still a lengthy process, and outside the scope of this work, but we believe it strikes a good balance between robustness and speed and has the potential to expedite drug discovery. CPB is a software tool designed to enable this approach, and we hope that by releasing it we engage more researchers in HCCT planning, leading to a gradual unraveling of its best practices and clinical efficacy.

## Current Limitations and Future Work

CPB makes use of TDM to surface relevant drugs and drug interactions, but there are number of important aspects it does not currently address, which are subject to future work:

1. CPB identifies drug combinations and prioritizes synergistic ones but does not explicitly identify and warn against adverse drug reactions.
2. CPB does not address the temporal aspects of treatment planning. While the graphical interface allows researchers to specify temporal notes on edges (e.g., “given 24 hours after” or “given together with”), identifying the relevant info in the scientific papers is left to the researcher.
3. Similarly, CPB does not currently suggest possible dosages for the drugs in a treatment plan. As before, this can be added manually as treatment notes, but identifying the relevant information in the literature is not automated.
4. CPB suggestions are limited to pairwise interactions (drug-disease and drug-drug), whereas higher order combinations with high synergy between components are not identified.

Each of the above issues can be addressed in the future by NLP modules integrated into CPB: e.g., AI models can be used to identify finer grained drug combinations (e.g., synergistic and antagonistic) along with temporal and dosage information. Moving from pairwise interactions to higher order combinations is another interesting line of research.

Despite these limitations, we show that the ‘interactive human in the loop’ approach manifested in CPB enables HCCT planning that is considerably faster and more comprehensive and evidence based than what is currently possible. We leave these additional aspects as areas for future research.

### Conclusion

To conclude, in this paper we develop a path to extend the boundaries of complexity in combinatorial cancer treatments. We first characterized the current complexity of combination therapies in cancer and showed that the current limit is 6 drugs per plan and the possible combinatorial space is huge. We then developed and evaluated a novel interactive computational tool based on TDM to generate reasonable plan of much larger sizes. We show that plans generated with the help of the tool in a few hours compare favorably to plans generated without it based on existing tools in the course of a month.

## Methods

### 1. Background: The SPIKE Extractive Search Engine

To perform text mining, we used the SPIKE extractive search engine^23,24^. SPIKE is a freely accessible search engine over PubMed which offers a number of relevant capabilities absent from PubMed Search (i) it allows to load very large lists of terms and use these in queries (compared to PubMed which limits searches to 2048 terms); (ii) it allows for fine grained control over the scope of the matches, e.g. requiring a treatment term and a cancer term to appear in the same sentence or even in a specific syntactic configuration; and (iii) it supports “extractive searches”, which means it not only finds sentences which match a query, but also extracts spans of text which correspond to designated query terms. For example, it not only returns sentences that match a drug from a list, but also extracts *which* drug was mentioned for each sentence; (iv) finally, it also allows a rich set of query types, including searching for sequential patterns and for sentence-structure patterns. We used SPIKE queries in our quantification of combination sizes in the literature, as well as to identify, aggregate and rank treatments in the CPB app.

### 2. Complexity Characterization

#### 2.1. Clinicaltrials.gov

We used clinicaltrials.gov database and downloaded the data of ~7,300 trials from the past 10 years to a CSV file (file downloaded in July 2021). The search was filtered for ‘cancer’ in the disease field and the phrases ‘combination’ or ‘combined’ in the intervention field. From the CSV file we analyzed the different interventional arms in each trial and extracted the number of elements used in each one. We also extracted the number of drugs from the titles of the trials.

#### 2.2. Text Data Mining (TDM) for Quantifying Combination Sizes

We used the SPIKE TDM engine in two stages. First, we searched for different phrasings used in published papers to describe drug combinations or drug regimens (for example, the phrasings ‘drug combination’, ‘drug Regiment’ or ‘combining five drugs’, etc.), this step was done by using the structure search in SPIKE search engine. Then, using the list formed in the first step and basic search, we extracted the number of drugs combined in a single treatment in all diseases, in cancer in general and in specific types of cancer. Searches were done in July 2021.

### 3. CPB Tool Development

#### 3.1. Searching for First Line Treatments

To support the search and extraction of treatments, we pre-curated a comprehensive list of drugs, and non-drug cancer treatments. The drug list was derived from drugbank.com with manual deletion of supplementary food, minerals and vitamins. Based on the input cancer name, genetic findings and additional keywords (hereafter referred to as biomarkers), the app runs a number of SPIKE-PubMed queries using a backoff strategy.Queries are run in succession until the 100 most related treatments are obtained.

- The first query is for sentences containing a treatment from the treatments list and a biomarker from the patients’ biomarkers. The sentences are restricted to originate from papers whose **title** includes the input cancer name
- If the result set does not include at least 100 unique treatments, we run a similar query. This time, restricting the result sentences to originate papers whose **abstract** contain the input cancer name. We add the additional results into the result set.
- If the result set still does not include 100 unique treatments, we run a similar query, this time allowing for biomarkers to appear in the same paragraph as the treatment (rather than the same sentence).

We extract the treatment from each of the result sentences and score a treatment by the number of matches which referenced it. To reflect the fact that the queries are ordered by the likelihood of retrieving relevant treatments, a treatment gets 3 points for every match due to query 1, 2 points for matches due to query 2 and 1 point for matches due to query 3. The treatments are then ranked by their aggregated score and displayed to the user. The evidence for each treatment is a list of abstracts where matching sentences are highlighted as well as the relevant treatment and matched biomarkers in them. The evidence abstracts are ranked by (i) the unique number of matched biomarkers in them, and (ii) in case of a tie, the total number of matched biomarkers in them. Note that the ranking system is a critical component of the solution and massively reduces planning time. Instead of reviewing hundreds of papers in order to identify treatments and sort which ones are more frequent and more related to the input conditions, the app immediately surfaces the specific treatments which are most frequently discussed in the literature. Also, the ranking within the evidence snippets of a treatment leads to planners typically needing to review only 1-3 specific sentences to validate the treatment relevance

#### 3.2. Searching for Synergistic Treatments

When the user clicks an existing treatment in the plan, the app searches PubMed for treatments which are potentially synergistic to it (with or without reference to the given cancer name). To support this search, we curated a list of terms whose presence in a sentence along with a pair of treatments typically indicates a synergistic combination. As before, we run a sequence of SPIKE-PubMed queries using a backoff strategy:

- The first query is for sentences containing the input treatment, another treatment from our treatments list and a term indicating a synergistic combination. The sentences are restricted to originate from papers whose **title** contains the input cancer name.
- If the result set does not include 100 unique treatments, we rerun the query, this time restricting the result sentences to papers whose **abstract** contains the input cancer name.
- If the result set still does not include 100 unique treatments, we rerun the query, this time **not restricting** the sentences to appear together with the input cancer name.

The ranking of treatments and of the evidence for each treatment is proceeds as described for first line treatments.

### 4. Integrating Supervised Machine Learning (ML) Models

Supervised ML models typically have higher coverage then pattern-based methods, but they also introduce some additional noise. Integrating such models into our app is straightforward. Given a model which identifies drug combinations in text and labels the combination as synergistic or antagonistic, extract all such combinations from PubMed into a database, along with confidence scores and frequency data. This database can then be queried by the planning app to suggest treatments to the user. Based on Tiktinski et al^25^, we also integrated the output of ML models for drug combinations to the app, in an optional mode that allows the planner to use the more precise and more replicable non-ML results, or to extend the results with the higher coverage but noisier ML output. The experiments in this work are conducted on the more precise mode

## Supplementary

**Supplementary Fig S1.**
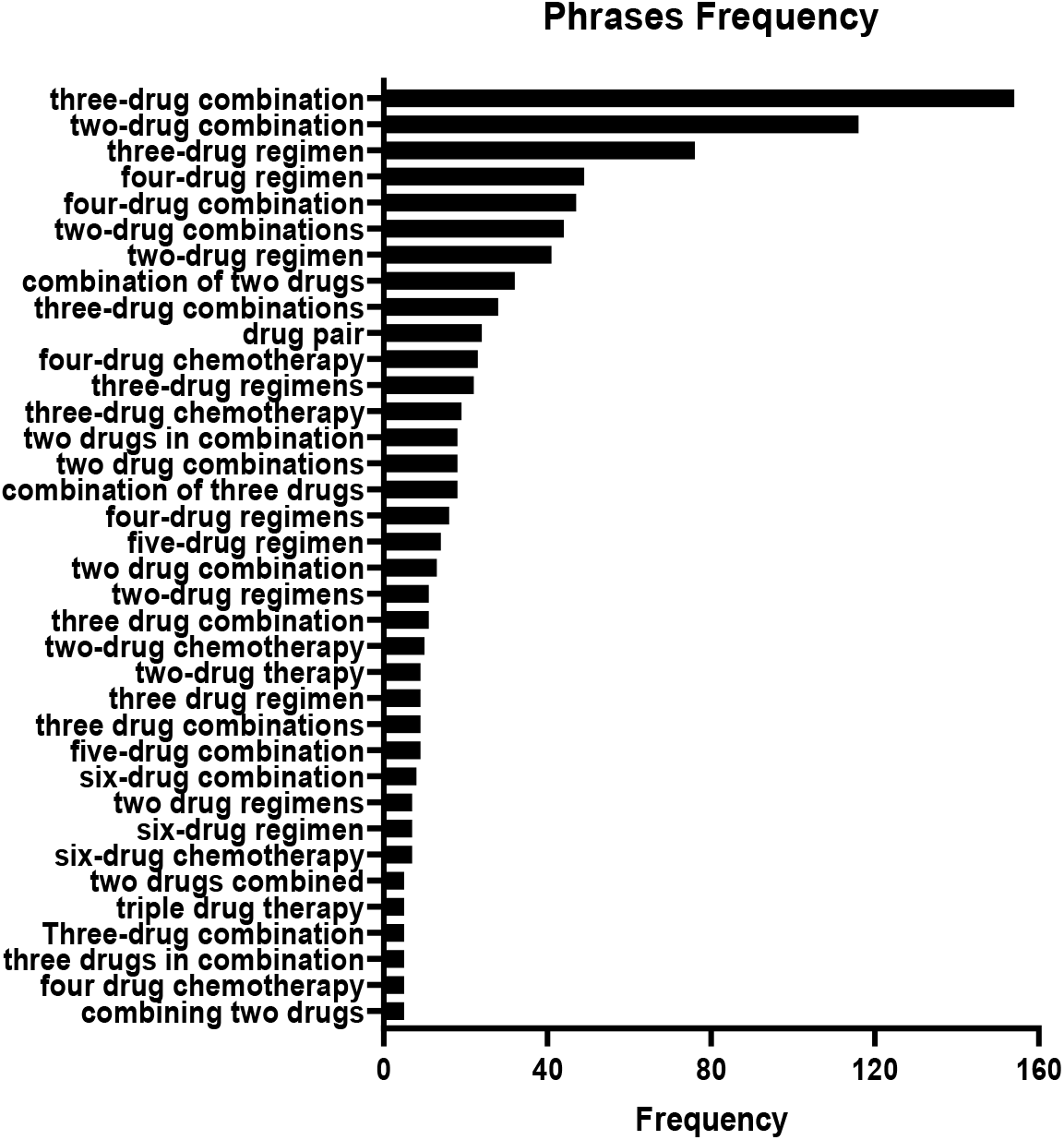
Frequency of phrasing combinatorial drug treatments.

**Supplementary Fig S2.**
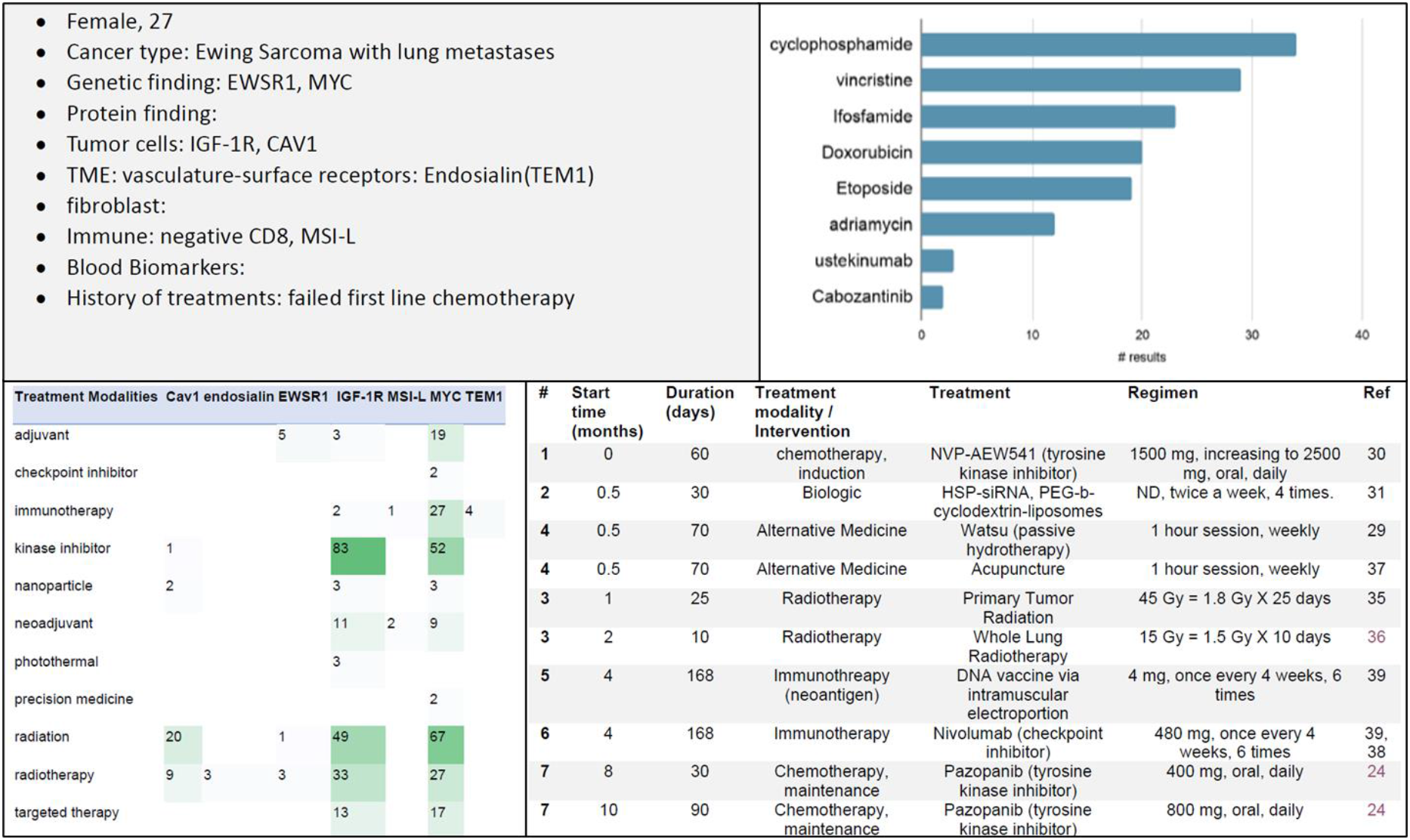
elements from an example stunent’s paper, decryptions from top left to bottom right. (A) the case study and patients characteristics, (B) relevant drugs for Ewing sarcoma (extracted from SPIKE), (C) the number of articles connecting between treatment modalities and the protein findings, (D) the final, detailed treatment plan.

